# Urbanization drives parallel adaptive clines in plant populations

**DOI:** 10.1101/050773

**Authors:** Ken A. Thompson, Marie Renaudin, Marc T.J. Johnson

## Abstract

Urban areas are a new and increasingly dominant feature of terrestrial landscapes that dramatically alter environments. It is unclear whether wild populations can adapt to the unique challenges presented by urbanization. To address this problem, we sampled the frequency of a Mendelian-inherited trait—cyanogenesis—in white clover (*Trifolium repens* L.) plants along urbanization gradients in four large cities. Cyanogenesis protects plants from herbivores, but also reduces freezing tolerance. Plants evolved reduced cyanogenesis with increasing proximity to the urban center in three of the four cities. In an experiment, we demonstrate that gradients in herbivore pressure do not cause these clines. Instead, urban areas experience relatively cold minimum winter ground temperatures because of reduced snow cover within cities, which selects against cyanogenesis. Together, our study demonstrates that wild populations exhibit parallel adaptive evolution in response to urbanization, which likely facilitates the persistence of these plants and promotes pollinator abundance and diversity.

## Introduction

Cities and urban environments are a novel and increasingly dominant feature of terrestrial landscapes where more than half of the world’s human population now reside (United Nations 2014). Relative to non-urban areas, urban environments have dramatically different environmental conditions, such as warmer air temperatures (Oke 1973) and altered ecological communities (Raupp et al. 2010). Whereas urbanization can have large ecological effects on plant and animal communities (Hahs et al. 2009), little is known about whether wild populations are adapting to urban environments (Cheptou et al. 2008; Donihue and Lambert 2014; Alberti 2015; Johnson et al. 2015; Munshi-South and Harris 2016). To investigate evolutionary processes in urban ecosystems, we used a globally distributed perennial plant, white clover (*Trifolium repens* L.), as a model to examine adaptive evolution along urbanization gradients.

White clover exhibits a Mendelian-inherited polymorphism for the production of hydrogen cyanide (HCN), a chemical defense against herbivores that inhibits cellular respiration and is toxic to both plants and animals. Within populations, cyanogenic genotypes produce HCN following tissue damage, whereas HCN is absent from acyanogenic genotypes (Hughes 1991). The cyanogenesis polymorphism is caused by two genes that each encode a chemical component of cyanogenesis: *CYP79D15* (hereafter *Ac*) and *Li;* Plants must have dominant alleles at both loci to produce HCN (Olsen et al. 2014). The two components are stored separately in cells to avoid self-toxicity, and form HCN when they are brought together following tissue damage. Previous research found that cyanogenesis is most frequent in populations at low latitudes and elevations, where herbivory is often highest and temperatures are warmer (Daday 1954a). It has been hypothesized that herbivores select for cyanogenesis in these warm environments (Hughes 1991; Kooyers and Olsen 2012). In cold climates, cyanogenesis is selected against because freezing temperatures lyse cells and trigger HCN release, causing self-toxicity in cyanogenic plants (Daday 1958; Hughes 1991).

Environmental change associated with urbanization may alter natural selection on cyanogenesis. Whereas herbivores are not consistently affected by urbanization (Raupp et al. 2010), air temperatures are generally warmer in urban areas relative to non-urban areas (Landsberg 1981). This may cause urban populations to experience fewer freezing events than non-urban populations during winter (Parris and Hazell 2005), and consequently weaken selection against cyanogenesis in cities. Thus, we predicted that cyanogenesis would increase in frequency with increasing proximity to urban centers. We surveyed 2379 plants from 121 populations of white clover along three 50-km transects radiating outward from downtown Toronto, Canada (Fig. S1), and screened them for their ability to produce HCN. To confirm that selection is acting on HCN, and not on *Ac* or *Li* individually for unrelated functions, we also quantified the frequencies of the *Ac* and *Li* alleles.

## Results and Discussion

We found consistent changes in the frequency of cyanogenesis along each transect, but clines were in the opposite direction to our prediction—cyanogenesis frequency decreased with increasing proximity to downtown Toronto at a rate of 0.65 ± 0.19% km^−1^ (*F*_1, 119_ = 30.57, *P* < 0.001) (Fig. 1A; Fig. S2). The frequency of dominant alleles at both *Ac* and *Li* also decreased toward the urban center, which is consistent with selection acting on cyanogenesis rather than *Ac* or *Li* individually (Kakes 1987) (Fig. S3). Local habitat variables like human population density and percent impervious surface cover did not explain significant variation (all *r*^2^ ≤ 0.06; all *P* > 0.6) in cyanogenesis frequency in multiple regression models that also included distance from the urban center. Collectively, these results suggest that large-scale biological and/or climatic processes associated with urbanization drive adaptive clines in cyanogenesis.

**Fig. 1.**
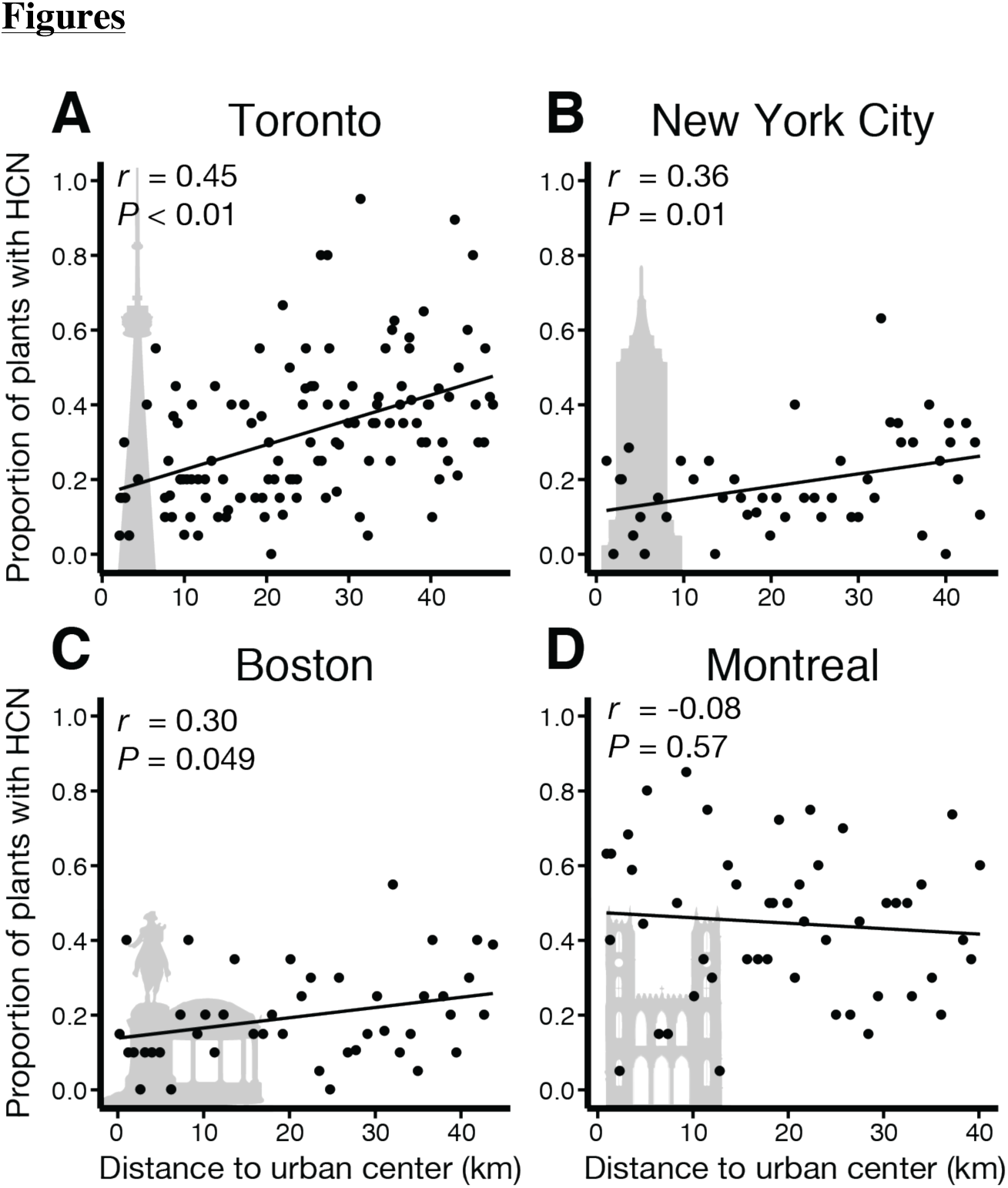
Urban populations of white clover, *Trifolium repens*, have evolved reduced cyanogenesis relative to non-urban populations in three cities. We observed parallel clines in cyanogenesis in: (**A**) Toronto, (**B**) New York City, and (**C**) Boston, but not in (**D**) Montreal. Each point represents the proportion of plants within a population that tested positive for cyanogenesis (mean *n* per population = 19.7 plants). Statistics were calculated using linear regression.

Parallel evolution to independent environmental clines provides some of the strongest evidence for evolution by natural selection (Schluter et al. 2004). To test for parallel evolution to urbanization gradients, we sampled white clover populations along one transect in each of Montreal (969 plants from 47 populations), New York City (946 plants from 48 populations), and Boston (876 plants from 43 populations) (Fig. S4–6). The frequency of cyanogenesis decreased toward the urban centers of both New York (0.34 ± 0.13% km^−1^; *F*_1, 46_ = 6.88, *P* = 0.01) and Boston (0.27 ± 0.14% km^−1^; *F*_1, 42_ = 4.09, *P* = 0.049), but there was no cline in Montreal (−0.14 ± 0.25% km^−1^; *F*_1, 47_ = 0.33, *P* = 0.57) (Fig. 1B–D). Because we did not observe a cline in Montreal, we hypothesized that selective mechanisms causing clines in the other cities are absent there. We conclude that adaptation in response to urbanization best explains the observed parallel clines in cyanogenesis across multiple cities.

Adaptive clines in cyanogenesis can arise because of spatial gradients in either the defensive benefits or the self-toxicity costs of cyanogenesis (Kooyers and Olsen 2012). We tested two non-exclusive hypotheses that could explain our results: (i) cyanogenesis is more beneficial in rural areas because of increased herbivory, and (ii) cyanogenesis is more costly in urban areas because of colder winter ground temperatures. Urban areas typically have warmer air temperatures than non-urban areas (i.e., urban heat islands), and so it is non-intuitive that urban ground temperatures might be cold relative to non-urban areas. However, relative to non-urban areas, cities typically experience less snowfall and more snowmelt because of their warmer temperatures (Landsberg 1981). Snow cover insulates the ground against freezing temperatures (Goodrich 1982), and thus we explored whether a reduction in urban snow cover may cause plants in urban areas to experience colder winter temperatures than plants in rural areas.

To test if the defensive benefits of cyanogenesis change along urbanization gradients, we conducted a field experiment examining spatial variation in herbivory. We established 40 experimental white clover populations, each with equal numbers of cyanogenic and acyanogenic genotypes, on lawns located between downtown Toronto and rural areas west of the city (Fig. S7). Cyanogenic plants received less herbivore damage than acyanogenic plants during the early (*F*_1, 45_ = 6.57, *P* = 0.01) and late (*F*_1, 45_ = 7.85, *P* < 0.01) growing season. However, there was no change in overall herbivory (*β* = −0.13 ± 0.13%, *F*_1, 36_ = 1.04, *P* = 0.31) or relative herbivory on cyanogenic plants compared to acyanogenic plants (*β* = −0.001 ± 0.001%, *F*_1, 36_ = 1.07; *P* = 0.31) along the urbanization gradient (Fig. 2). Thus, the defensive benefits of cyanogenesis are unlikely to explain the observed clines.

**Fig. 2.**
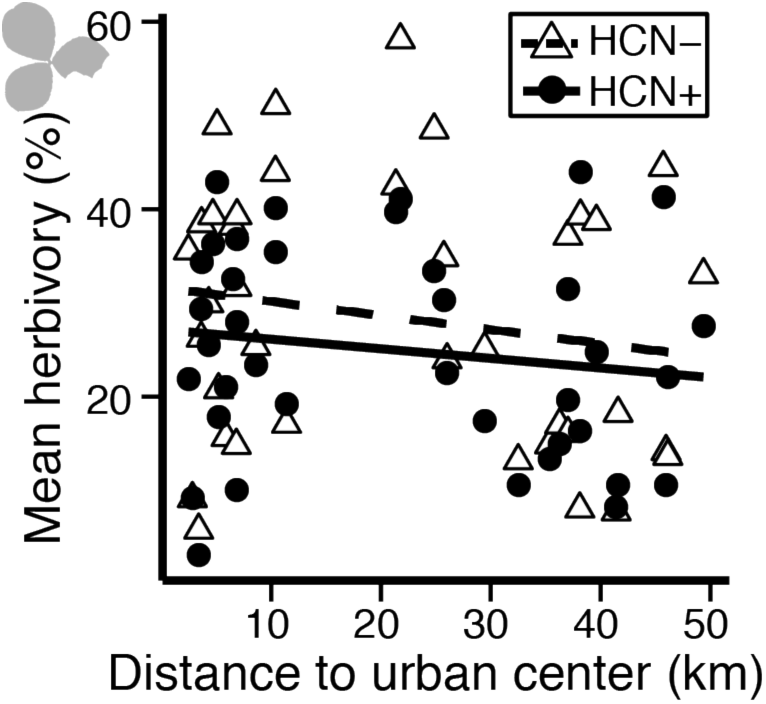
The benefits of cyanogenesis as a defense against herbivores do not explain the observed clines. Our field experiment revealed that cyanogenic plants (HCN+) received less leaf herbivory than acyanogenic plants (HCN-) overall (*P* < 0.05, ANOVA), but mean herbivory on both cyanotypes within experimental populations (each *n* = 38) changed little along an urbanization gradient in Toronto (both *P* > 0.1, ANOVA). Damage scores are the mean of early and late season herbivory.

To test whether clines in cyanogenesis could be caused by spatial variation in winter freezing temperatures, we placed temperature probes along an urbanization gradient in Toronto during the winter of 2015 (Fig. S8). We found that minimum ground surface temperatures were often coldest in urban areas (Fig. 3A). Using data for regional snow depth during this period, we found that ground temperatures in urban areas were colder than in non-urban areas when there was deep snow in the region, and warmer when there was little or no snow in the region (*F*_157_ = 32.1; *r*^2^ = 0.44, *P* < 0.01) (Fig. 3B). High regional snow cover likely provides the necessary conditions for clines in snow depth—and thus ground temperature—to arise along urbanization gradients. To test whether the white clover populations we sampled in the field occur along a snow cover gradient, we used remote sensing to quantify the normalized-difference snow index (NDSI) (Shimamura et al. 2006) for our sampled populations. We found that NDSI decreased (i.e., less snow) toward the urban center in each city (*β* = 3.70 × 10^−3^ ± 1.02 × 10^−3^, *F*_1, 228_ = 30.03; *P* < 0.01) (Fig. 4A). These observations collectively suggest that clines in winter minimum ground temperatures along urbanization gradients cause clines in cyanogenesis due to higher self-toxicity in urban habitats.

**Fig. 3.**
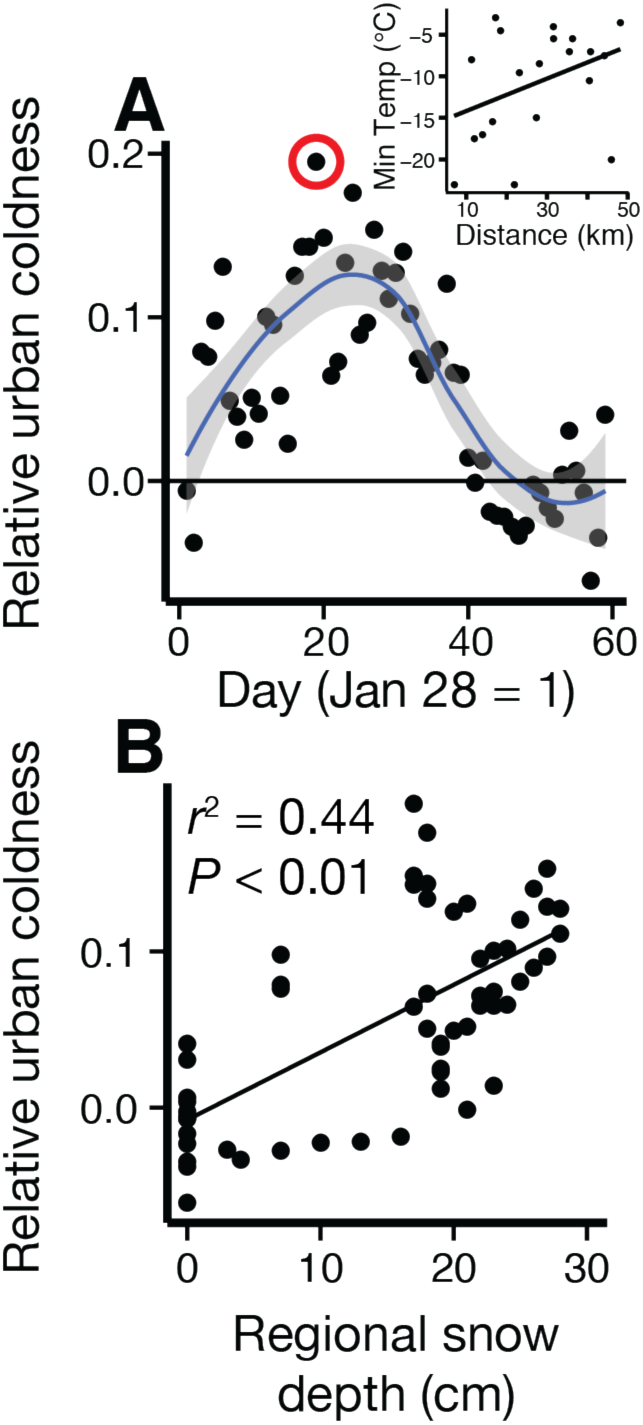
Urban ground temperatures are often colder than non-urban ground temperatures. (**A**) Analysis of daily relative urban coldness values reveal that it was often colder in Toronto during winter 2015 than in non-urban areas (curve is a 95%-CI loess-smoothed surface). We hypothesized that relatively cold urban ground temperatures are caused by urban-rural snow depth gradients, and consistent with this expectation we found (**B**) a significant positive correlation between the daily relative urban coldness index value and regional snow depth on those days.

We sought to explain why clines in cyanogenesis occur in Toronto, New York City, and Boston, but not Montreal. Because insulation provided by snow cover could weaken selection against cyanogenesis, we predicted that urban plants are most insulated by snow cover in Montreal. We found that mean NDSI was significantly higher in Montreal (i.e., more snow) than all other cities (pairwise contrasts, all *P* < 0.02), which did not differ from each other in mean NDSI (pairwise contrasts, all *P* > 0.5) (Fig. 4A). Using weather station data from each city, we calculated daily snow depth and minimum temperature between 1980 and 2005. We found that Toronto, Boston, and New York City have more days with < 0 °C temperatures and no snow cover than Montreal (Tukey HSD, all *P* < 0.01) (Fig. 4B). On < 0 °C days, snow is > 5 cm deeper in Montreal than the other three cities (Tukey HSD, all *P* < 0.01) (Fig. 4C). Moreover, cyanogenesis frequency is highest in Montreal (Fig. S9). These results are consistent with greater snow cover in Montreal protecting plants from freezing and relaxing selection against cyanogenesis.

**Fig. 4.**
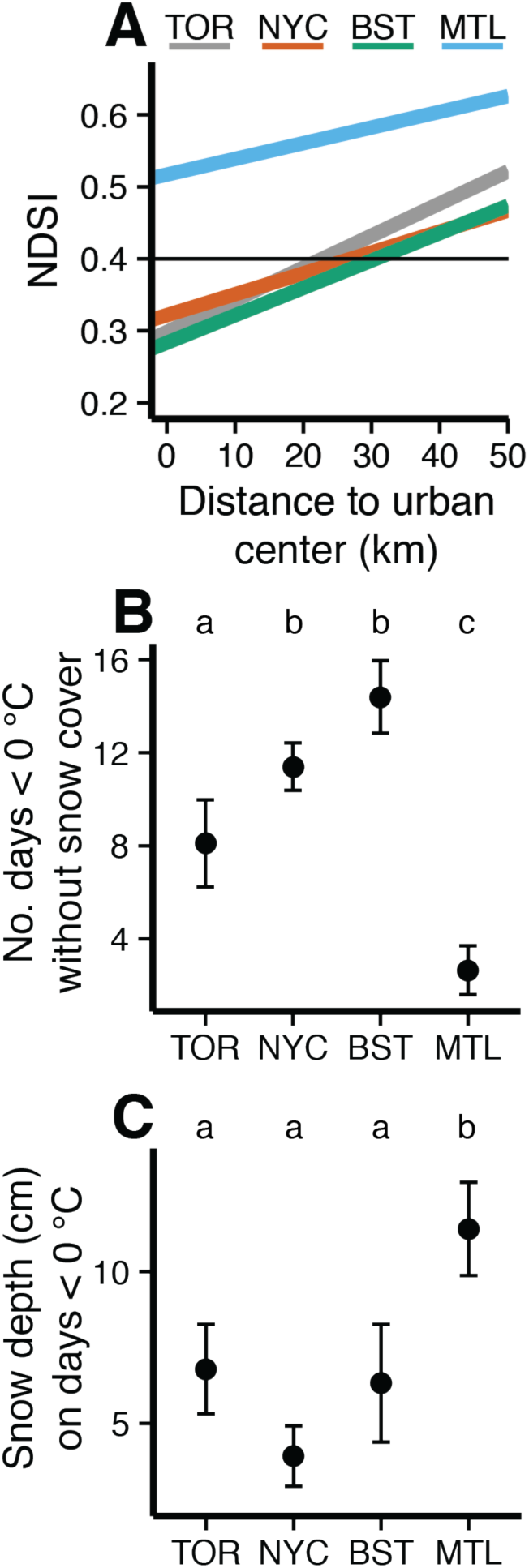
Historic differences in January-February snow cover and temperature across the four cities sampled in this study. (**A**) The normalized-difference snow index (NDSI) decreased toward the urban center of each city. The horizontal line at NDSI = 0.4 represents the threshold value for the presence of snow (Shimamura et al. 2006). (**B**) Montreal (MTL) has fewer < 0 °C temperature days without snow cover (± 1 SE), and (**C**) deeper snow cover (± 1 SE) on days where temperatures are < 0 °C, than Toronto (TOR), New York City (NYC), and Boston (BST). Different letters represent significant differences at *P* < 0.05 (Tukey HSD).

Our results suggest that urban-rural gradients in winter ground temperature, which arise due to snow cover gradients, drive parallel adaptive clines in cyanogenesis. These conclusions are consistent with several studies of cyanogenesis in *T. repens* that experimentally demonstrate reduced cyanogenesis as an adaptation for freezing tolerance (Daday 1965; Dirzo and Harper 1982b; Caradus and Eerens 1989; Hughes 1991; Olsen and Ungerer 2008), and hypothesize that high snow cover may prevent the evolution of clines in cyanogenesis (Till-Bottraud and Gouyon 1992). Genetic variation for cyanogenesis is likely maintained by a balance between gene flow and spatial variation in selection (Supplementary Text S2-3).

Although adaptation to urban environments can reduce the risk of extinction in cities, it is currently unclear whether wild populations are adapting in cities (Cheptou et al. 2008; Donihue and Lambert 2014; Johnson et al. 2015; Munshi-South and Harris 2016). Our study demonstrates that wild plant populations are capable of rapidly adapting to the unique environmental conditions caused by urbanization. White clover is the most abundant source of nectar for pollinators in many temperate ecosystems (Baude et al. 2016). Thus, the ability of white clover to rapidly adapt to urban ecosystems may not only contribute to its own dominance in cities, but could also facilitate diverse and abundant pollinator communities in urban ecosystems.

## Methods

### Surveys of natural populations

#### Field sampling

We sampled 128 populations (2509 plants) of *T. repens* along three transects in the Greater Toronto Area (GTA) (Fig. S1). Transects extended west, north, and east of Toronto. For both the east and west transects, we attempted to maintain a constant distance from Lake Ontario. We stopped approximately every 1 km to sample populations of *T. repens* along each transect. For each population, we recorded GPS coordinates and elevation, and collected stolon cuttings from up to 20 plants; cuttings were approximately 4 cm in length and had at least two leaves. To reduce the likelihood of sampling a single genotype twice, we ensured that there was at least 1.5 m between all collected stolons. We placed all stolons from a population in a sealed bag, and stored them in a cooler with ice. At the lab, we put one leaf from each plant into wells of 96-well plates and stored the rest of the tissue in microcentrifuge tubes. We stored plates and tubes at −80 °C. Seven populations had coarse netting above the soil and were excluded from the analysis because they may have been seeded in recent years as a soil stabilizer; our conclusions do not change when these populations are included in the analysis. To determine if parallel clines in cyanogenesis occur in other cities, we sampled one 50-km transect in each of Montreal, QC, Boston, MA, and New York City, NY, between 26 July and 1 Aug 2015 (Fig. S4–6). Sampling and tissue storage methods were identical to those outlined above.

#### Cyanogenesis analysis

To determine the frequency of cyanogenesis in natural populations of *T. repens*, we tested each plant for the presence of HCN using Feigl-Anger assays. Feigl-Anger assays determine the presence or absence of HCN in a sample using a simple colour change reaction (Thompson and Johnson 2016). We removed the 96-well plates that contained leaves from the freezer, allowed them to thaw, and added 80 *μ*L of water to each well. We used pipette tips to macerate the tissue in the wells, and secured Feigl-Anger test paper over the plates, taking care not to let any water or tissue contact the paper. We incubated the plates for 3 hr at 37 °C, and then scored each well for cyanide (i.e., cyanotype *AcLi*), which is indicated by a blue colour. We recorded the proportion of plants that were cyanogenic in each population.

To confirm that clines were due to selection on HCN and not *Ac* or *Li*, we screened the frequency of *Ac* and *Li* from acyanogenic plants collected from the three Toronto transects. There was insufficient leaf tissue to screen every plant at both loci, so we screened all acyanogenic plants in a particular population for either *Ac* or *Li*, alternating the locus assayed with every sequential population. Screening was conducted as above but instead of 80 *μ*L of H_2_O, we added either 30 *μ*L of 10mM linamarin (Sigma-Aldrich 68264) and 50 *μ*L H_2_O to each well to screen for linamarase (cyanotype *acLi*), or 80 *μ*L of 0.2 EU/mL linamarase (LGC Standards CDX-00012238-100) with 20 *μ*L of H_2_O to each well to screen for linamarin (cyanotype *Acli*). Our earlier study confirmed that Feigl-Anger assays correspond with the presence/absence of the *Ac* and *Li* gene sequences (Thompson and Johnson 2016).

#### Statistical analysis

We used linear regression to determine if the frequency of cyanogenesis was associated with urbanization. We first used the haversine formula—which calculates the distance in kilometers between two sets of co-ordinate—to determine the distance from each sampled population to the urban center (Table S1):

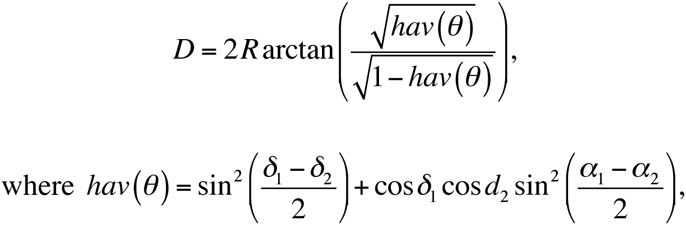

(latitude, longitude) co-ordinates of the two points are (*α*_1_, *δ*_1_) and (*α*_2_, *δ*_2_) in radians, and *R* is the radius of earth (6,371 km). Next, we regressed the frequency of cyanogenesis in populations against the distance of those populations from the urban center. All models had the proportion of plants in a population that tested positive for HCN as the response variable and distance to the urban center as a predictor. All statistical analyses were conducted using R 3.0.3 (R Core Team 2014).

While distance to an urban center might explain adaptive patterns, it is important to distinguish large-scale factors associated with urbanization from small-scale local factors. To accomplish this, we retrieved 2015 population density estimates from the NASA GPW4 dataset (resolution = 1 km) (CIESIN 2015). We also retrieved data for percent impervious surface cover (resolution = 30 m) (https://coast.noaa.gov/dataregistry/search/collection/info/nlcd-impervious), which has recently been shown to predict evolutionary patterns in urban environments (Munshi-South and Harris 2016). To determine if these variables explained variation in the frequency of cyanogenesis, we fit generalized linear mixed effects models as before, but added a term for population density (all cities) or percent surface imperviousness (New York City and Boston only). We found that in all cities, population density and percent surface imperviousness did not significantly explain variation in cyanogenesis when distance to the urban center was included in the model (all *r*^2^ < 0.06; *P* > 0.6). On its own, population density was a significant predictor of cyanogenesis frequency in Toronto and New York City, but in both cases its explanatory power (*r*^2^) was less than half of that of distance to the urban center. Percent impervious surface cover did not significantly predict cyanogenesis frequency. Thus, we concluded that large-scale factors associated with urbanization are more important for driving the evolution of cyanogenesis clines than fine-scale local factors.

### Field Experiment

#### Experimental design

We conducted a field experiment in the Greater Toronto Area (GTA) to test if herbivory is associated with urbanization, and if this could explain clines in cyanogenesis along urbanization gradients. We created forty small populations of *T. repens* on private lawns that spanned from downtown Toronto to suburban and rural areas west of the city (Fig. S7). The strongest individual cline in cyanogenesis was observed along this transect (Fig. S2A). From 4–6 June 2015, we placed 24 *T. repens* plants into solid plastic flats on a 1.5 m^2^ area of each lawn. We used solid flats to allow rainwater to collect below the plants and to prevent plants from accessing the belowground environment. Trays had spouts 1.5 cm above the base. During dry periods we watered all populations. Two populations were destroyed during the first month of the experiment and excluded from analysis.

In the field, each population contained twelve cyanogenic (*AcLi*) and twelve acyanogenic (4 *Acli*, 4 *acLi*, 4 *acli*) plants. Plants were initially propagated from stolons sourced from a group of 48 genotypes with cyanotypes determined using methods outlined above, except plants were screened at least twice for HCN, or *Ac* and *Li* gene products. We propagated stolons that were 5 cm long with 2-5 leaves into 6” pots (AZE06000, ITML, Brantford, ON) filled with pro-mix potting soil (Sunshine Mix #1; Sun Gro Horticulture, Agawam, MA). Plants were saturated with water and kept in a growth chamber set to 15.3 h of 500 *μ*mol light, with a 25:20 °C day:night temperature cycle. Approximately 0.5 g of slow-release fertilizer (Nutricote Total 13-13-13 With Minor Nutrients, Florikan, Sarasota, FL) was added to each pot. We randomly assigned plants to populations such that: (i) each genotype did not occur more than once in a population, (ii) each genotype occurred in 20 of 40 populations, and (iii) each population satisfied the cyanotype ratio described above.

We estimated herbivore damage on each plant twice during the experiment, early (July 9-13) and late (August 19-23) in the growing season. To measure herbivory, we selected 5 of the oldest non-senesced leaves from each plant and visually quantified the proportion of the leaf surface area eaten. Previous field experiments with *T. repens* determined that this method is sufficient to demonstrate differences in herbivory between cyanogenic and acyanogenic plants (Thompson and Johnson 2016). During each visit, we collected all ripe inflorescences. Halfway through the experiment, we harvested the aboveground biomass of each plant. This harvesting mimicked the intermittent mowing that plants typically experience on lawns, and prevented plants from rooting in adjacent pots. We collected all inflorescences that were clipped in this process. We dried harvested biomass for 72 hr at 70 °C and weighed it to the nearest 0.1 g.

We harvested all plants in late August. We recorded the total lifetime number of inflorescences that each plant produced, and weighed the floral and vegetative tissue separately. After weighing and counting the inflorescences, we ground them through 1 mm and 0.5 mm sieves in series to isolate the seeds, which were weighed to estimate maternal fitness.

#### Statistical analysis

To assess whether cyanogenesis reduced herbivory, we conducted a linear mixed-effects model with HCN as a fixed effect factor, and population and genotype as discrete random effects. We fit separate models for both the early-and late-season surveys, using the average damage recorded across all measured leaves as a response variable. The full statistical model used in our analyses was:

Herbivory = intercept + HCN + genotype + population + error

To ensure that differences in herbivory were due to HCN and not *Ac, Li*, or linked genes, we fit separate models for a dataset that consisted of acyanogenic plants. These models were similar to those described above but instead of HCN, the predictor variable was the presence or absence of functional *Ac* or *Li* alleles.

To determine if herbivory and selection on cyanogenesis changed across the urbanization gradient, we calculated the mean herbivory in each population, and both relative fitness and relative herbivory for each plant within its own population. We used both lifetime and final vegetative biomass, and lifetime seed biomass as fitness components. To determine if these variables changed along an urbanization gradient, we regressed the population-level estimates of mean herbivory, and relative fitness and herbivory, against the distance of each population from the urban center of Toronto. To determine if herbivory and selection on cyanogenesis changed across the gradient, we took the mean value of relative herbivory, and relative fitness, for all cyanogenic plants in a population and regressed it against distance from the urban center. These values represent the relative fitness (or relative herbivory) of the *AcLi* genotype in a given population. Models all took the same form; as an example, the full statistical model for relative fitness of cyanogenic plants was:

*w*_i(HCN)_ = intercept + distance to urban center (km) + error

### Ground temperature data

During Winter 2015, we recorded ground temperature data across an east–west transect in the GTA (Fig S8). On 27 Jan 2015, we placed 20 iButton DS1921G thermochron devices (Maxim Integrated^™^, San Jose, USA), housed inside sandwich bags, at twenty locations approximately 5 km apart. We placed the iButtons underneath trees and bushes so they were not directly exposed to sunlight. We retrieved all iButtons on 28 March 2015. We extracted daily values for mean, maximum and minimum temperature, and standard deviation of temperature.

We used the temperature data to calculate a daily index of relative urban coldness. This index was calculated as the slope of minimum daily temperature versus the distance from the urban center of Toronto (Fig. S9). Positive index values indicate that minimum temperatures were colder in urban areas relative to non-urban areas, whereas negative values indicate that minimum temperatures were warmer in urban areas relative to non-urban areas. We treated each index value (*n* = 58) as an independent observation and fit a 95% CI locally weighted scatterplot smoothing function to visually evaluate if temperatures were significantly colder or warmer in the city during winter (Fig. 3A).

Because we observed that urban areas were often colder than non-urban areas, we hypothesized that this effect was due to differences in snow cover between the downtown core and outside non-urban areas. We retrieved snow depth data for the central Toronto weather station (WMOID: 71508; Lat: 43.6667, Long: −79.4000, Elevation: 113 m). This weather station is located approximately 2 km from our urban center reference point. We used a simple linear model with the relative urban coldness index value as the response and snow cover on the same day as the predictor. We found that snow cover was highly correlated with the relative urban coldness (Fig. 3B) and to a lesser extent maximum temperature and the standard deviation of daily temperatures (data not shown). The conclusions do not change when index values from all days are used.

### Landsat Data

To assess whether our study populations occurred along a gradient of snow cover, we used LandSat satellite data collected by the United States Geological Survey (USGS; earthexplorer.usgs.gov). We selected all images that met the following criteria: (i) image captured in January or February, (ii) no cloud cover above study area; (iii) images captured between 1980 and 2015; (iv) snow present along the transect (maximum NDSI ≥ 0.4; see below); (v) < 10 cm snowfall in the previous five days to allow any effects of urbanization on snow melt and/or removal to accumulate. We focused our analyses on January and February because January isotherms have previously been implicated as major factors driving the evolution of clines in *T. repens* cyanogenesis (Daday 1954b; Kooyers and Olsen 2012), and there were insufficient images from January alone for robust analyses. We note that the general pattern of decreasing NDSI toward urban centers is statistically significant when data from January alone are analyzed (*P* < 0.01).

For images that met these criteria (list will be archived online), we extracted the normalized-difference snow index (NDSI) for each of our study populations using the point sampling tool in QGIS 2.12. NDSI is a quantitative measurement of snow cover derived using the unique reflective and absorptive properties of snow in the green and short wave infrared (SWIR) wavelengths. It is calculated as (Green – SWIR) / (Green + SWIR). These wavelengths correspond to bands 2 and 5 for the Landsat 4, 5 and 7 satellites, and bands 3 and 6 for the Landsat 8 satellite. We maintained the standard 30 m cell resolution throughout our analysis because any resampling would bias urban areas toward lower NDSI due to a high density of artificial and paved surfaces, and bias rural areas toward higher NDSI for the opposite reason. Although Landsat data is at a resolution of 30 m × 30 m, we often sampled populations that were within 30 m of roads and were much smaller than the Landsat 30 m cells—especially in urban areas. Thus, using the GPS co-ordinates of populations would often cause us to include roads, sidewalks, or buildings in our analysis, which could bias our data toward detecting low NDSI values in cities. Because of this issue, we manually moved all of our points to the nearest large park or field before extracting NDSI data. Parks were larger than the NDSI grid cell size. A statistically significant, though weaker, associations remain between NDSI and urbanization remains when original locations are used.

We used linear mixed-effects models to calculate the change in snow cover vs. distance of populations to the urban center. We fit the following model to the data:

NDSI = intercept + distance + city + distance × city + month/year + population + image + error

where distance (from the urban center) is a continuous independent variable, city is a categorical fixed effect, month and year as nested random effects, and population (sampling location) and image both as categorical random effects. We analyzed the model using ANOVA with type-III sums of squares and the Kenward-Roger denominator degrees of freedom approximation to test the significance of fixed effects. We fit separate models for each city to extract the slope of NDSI vs. distance, and the y-intercept at distance = 0 km. These lines are plotted in Fig. 4A of the main text. To determine if mean NDSI was significantly different across cities, we fit the same model four times, but with the City ‘reference’ level changed across models. That is, one model compared Montreal to other cities, the next compared Toronto to other cities, *etc*. Because we had a small number of Landsat images in Toronto (*n* = 16), New York (*n* = 19) and Boston (*n* = 26), we did not correct for multiple testing when comparing mean NDSI across cities. When we conduct a Tukey HSD test, mean NDSI in Montreal is significantly different from the other three cities at *P* < 0.06, while mean NDSI does not differ between the other three cities (*P* > 0.9).

### Weather Station Data

To compare snow cover and snowfall across cities, we used data from weather stations. All data were retrieved through the National Climatic Data Center, which is operated by the US National Oceanic and Atmospheric Administration (NOAA; www.ncdc.noaa.gov/cdo-web/datatools/normals). We retrieved data for daily snow depth, snowfall, maximum temperature, and minimum temperature for the period of 1980 – 2010. We retrieved data from one airport weather station per city: Pierre Trudeau International Airport in Montreal, Lester B. Pearson International airport in Toronto, Logan International Airport in Boston, and Laguardia Airport in New York City. We excluded dates where these data were not available for all four cities and those that were flagged for quality. We treated all ‘trace’ values as zeroes.

After data were filtered, we had data for 3545 days in each city, which represented 20 years (1980–1992, 1994, 1996, 1999–2002, 2004). To test the hypothesis that plants in the four cities would differ in their exposure to snow cover, we calculated the cumulative number of days in January and February where temperatures were < 0 °C and there was no snow cover. This metric is strongly correlated with mean cyanogenesis frequency across the four cities (*t*_2_ = 6.74, *r* = 0.97, *P* = 0.02). We recorded this value for each year. Because the depth of snow influences its insulation properties (Goodrich 1982), we also compared the depth of snow on days where temperatures were < 0 °C.

Lastly, we compared snowfall between the cities, and found that Montreal has higher snowfall than the other three cities (Tukey HSD, all *P* < 0.01), which do not differ from one another (Tukey HSD, all *P* > 0.2). All models were simple mixed effects models that had the following form:

*Snow variable = intercept + city + year + error*

where city was a categorical fixed effect and year was a random effect.

## Acknowledgements

Thanks to C. Cameron, M. Escobar, M. Ganguli, R. Mittal, C. Prashad, J. Santangelo, and M. Urquhart-Cronish for help in the lab and field, and to many people who donated lawn space for our study (Supplementary Text S4). We thank A. Agrawal, S. Barrett, Y. He, P. Kotanen, J. Losos, H. Saini, D. Schluter, and the EvoEco lab for providing comments that improved the manuscript. KAT was supported by an NSERC CGS-M award and a Queen Elizabeth II Scholarship in Science & Technology. The project was supported by an NSERC Discovery Grant to MTJJ.

## Author contributions

K.A.T. and M.T.J.J. designed the study. All authors conducted fieldwork. K.A.T. conducted lab work and remote sensing analysis. K.A.T. analyzed the data with input from M.T.J.J. K.A.T. wrote the first draft of the manuscript, and K.A.T. and M.T.J.J. jointly revised the manuscript.

## Supplementary Figures

**Fig. S1.**
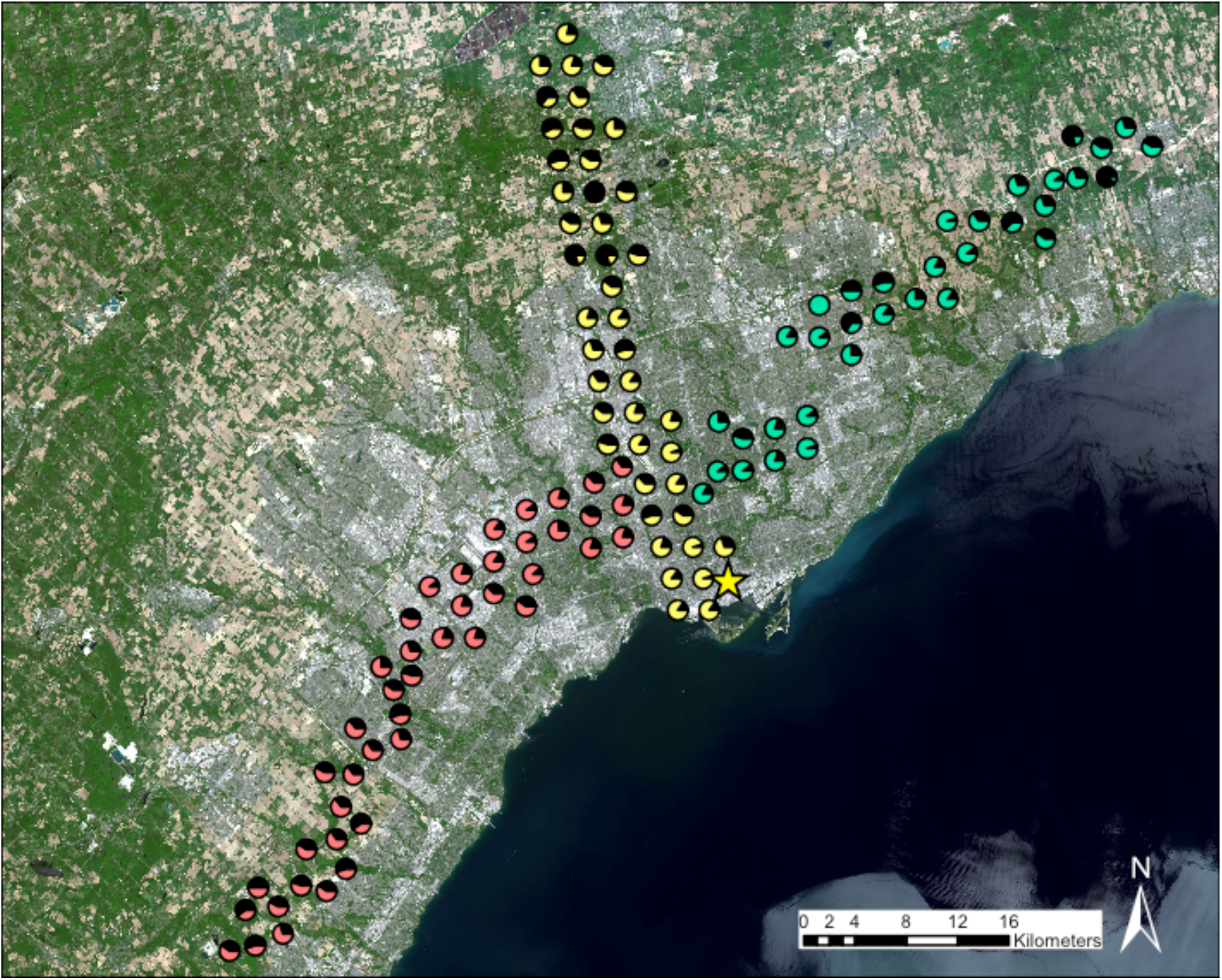
Map showing the location and cyanogenesis frequency of populations sampled in Toronto along transects that extended west (pink and black circles, *n* = 44), north (yellow and black circles, *n* = 43) and east (blue and black circles, *n* = 34) of the city center (total *n* = 121). The proportion of black colouration in pie charts reflects the frequency of cyanogenesis in that population. Pie chart locations are jittered to avoid overlap. We defined the urban center of Toronto as Yonge-Dundas Square (star symbol). The image is from the Landsat 8 satellite and was captured on 31 May 2014.

**Fig. S2.**
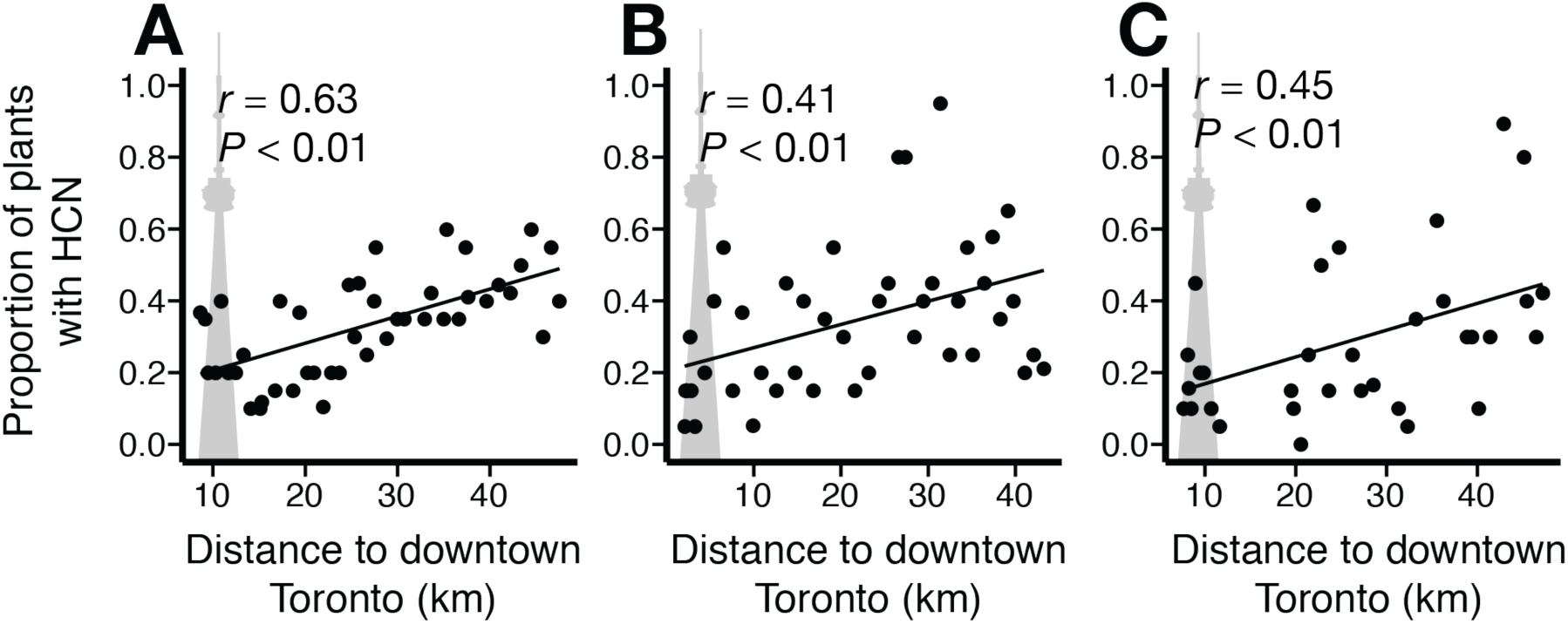
Regression of HCN frequency in natural populations of *Trifolium repens* along rural-urban transects extending: (**A**) west (*n* plants = 856, *n* populations = 44), (**B**) north (*n* plants = 843, *n* populations = 43), and (**C**) east (*n* plants = 680; *n* populations = 4 of downtown Toronto, Canada. We observed that the frequency of cyanogenesis decreased toward the urban center along each transect. Statistics were calculated using linear regression.

**Fig. S3.**
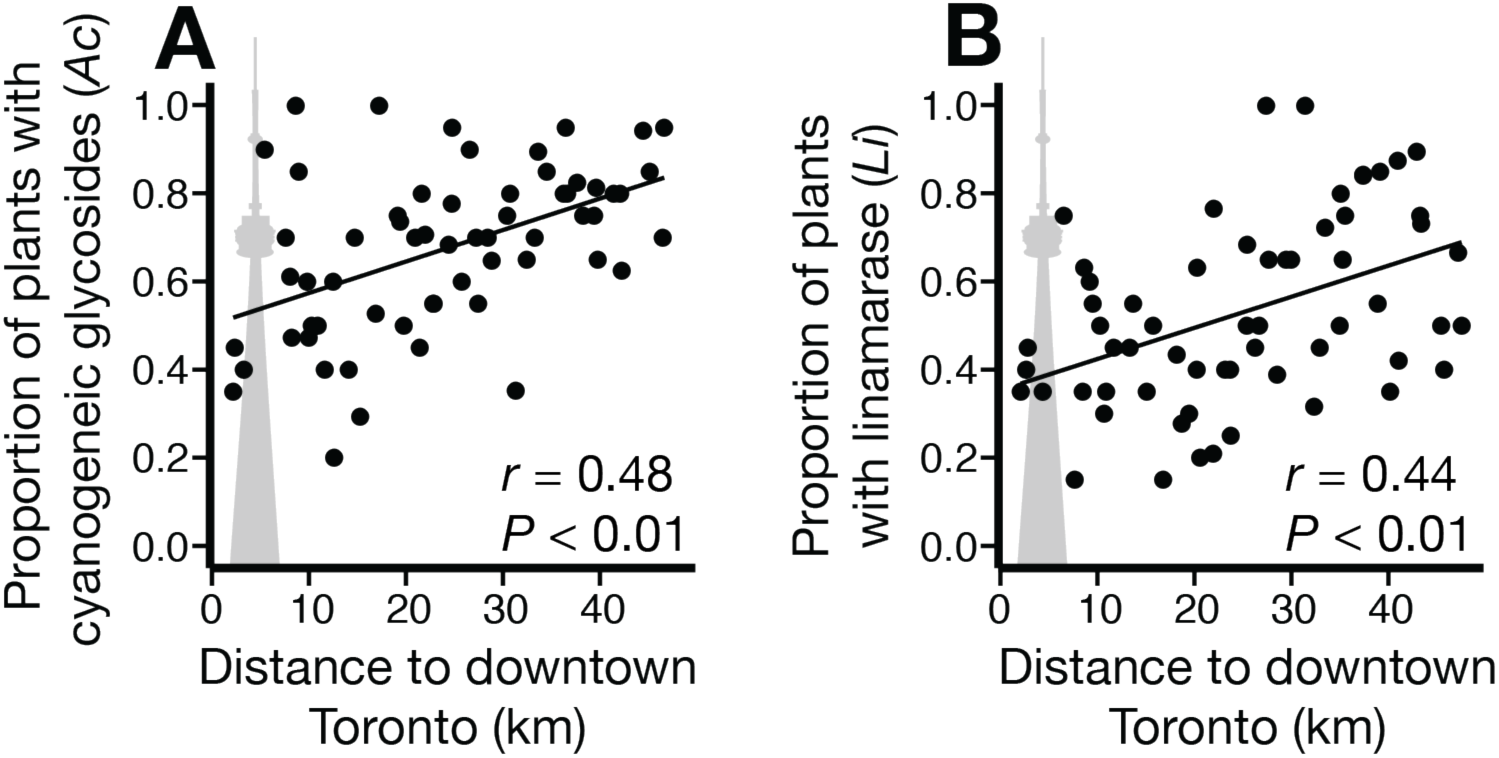
The frequency of functional alleles at: (**A**) *CYP79D15* (*Ac*) (0.71 ± 0.17% km^−1^; *χ*^2^_1_ = 9.45, *P* < 0.01) and (**B**) *Li* (0.71 ± 0.19% km^−1^; *χ*^2^_1_ = 12.00, *P* < 0.01) increase toward the urban center of Toronto. The dominant allele *Ac* encodes the primary step in cyanogenic glycoside biosynthesis, and the dominant allele at *Li* encodes the enzyme, linamarase. Data are pooled across all three transects. These results are consistent with the hypothesis that the epistatic interaction of the two loci (i.e., cyanogenesis) is responsible for urban-rural cyanogenesis clines, rather than either gene individually or linked genes. Statistics were calculated using linear regression.

**Fig. S4.**
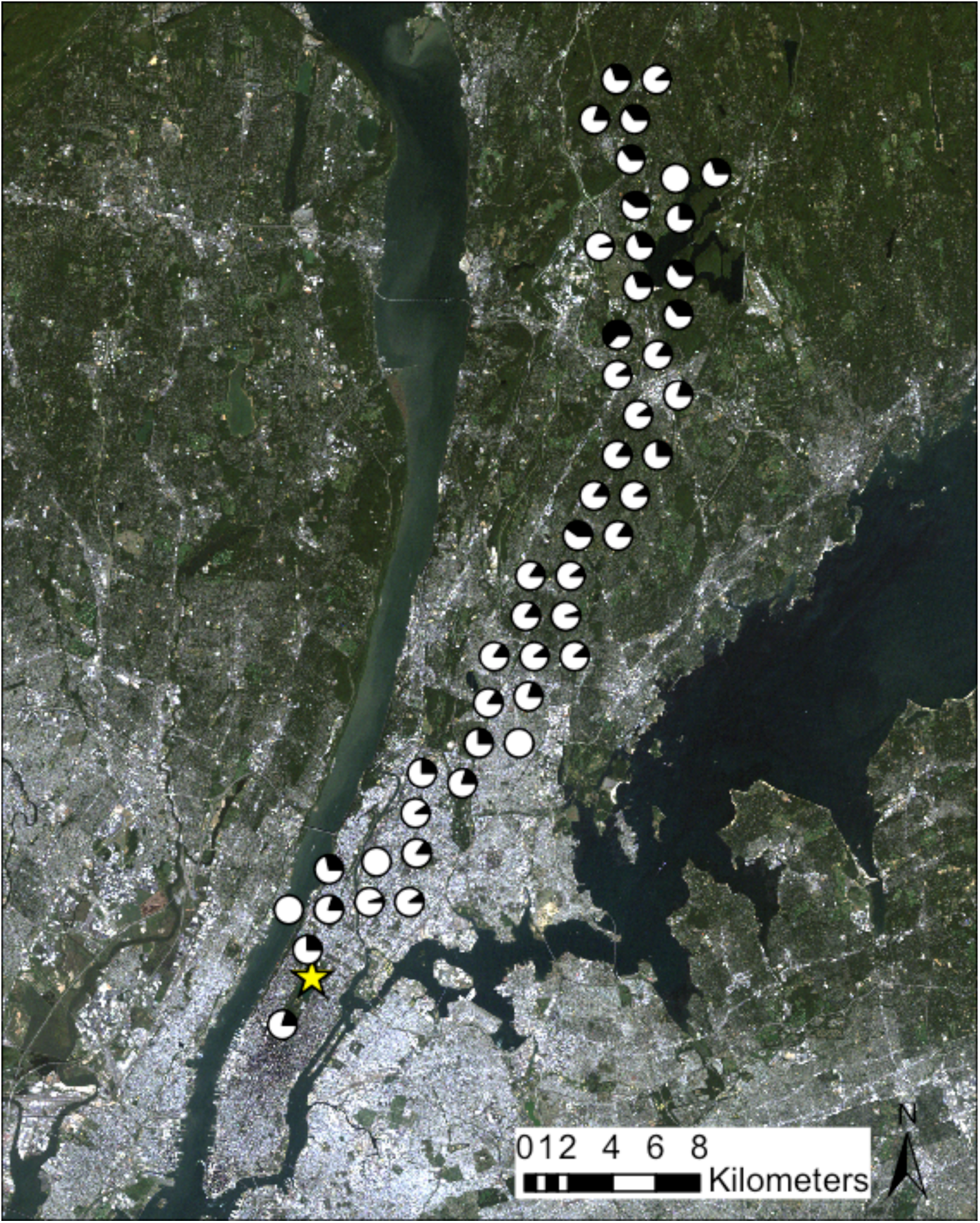
Map depicting the location and HCN frequency in populations sampled in New York City, USA (*n* = 48). The proportion of black colouration in pie charts reflects the frequency of HCN in that population. Pie chart locations were jittered to avoid overlap. We defined the urban center of New York as the northern border of Central Park (star symbol). The image is from the Landsat 5 satellite and was captured on 22 September 2010.

**Fig. S5.**
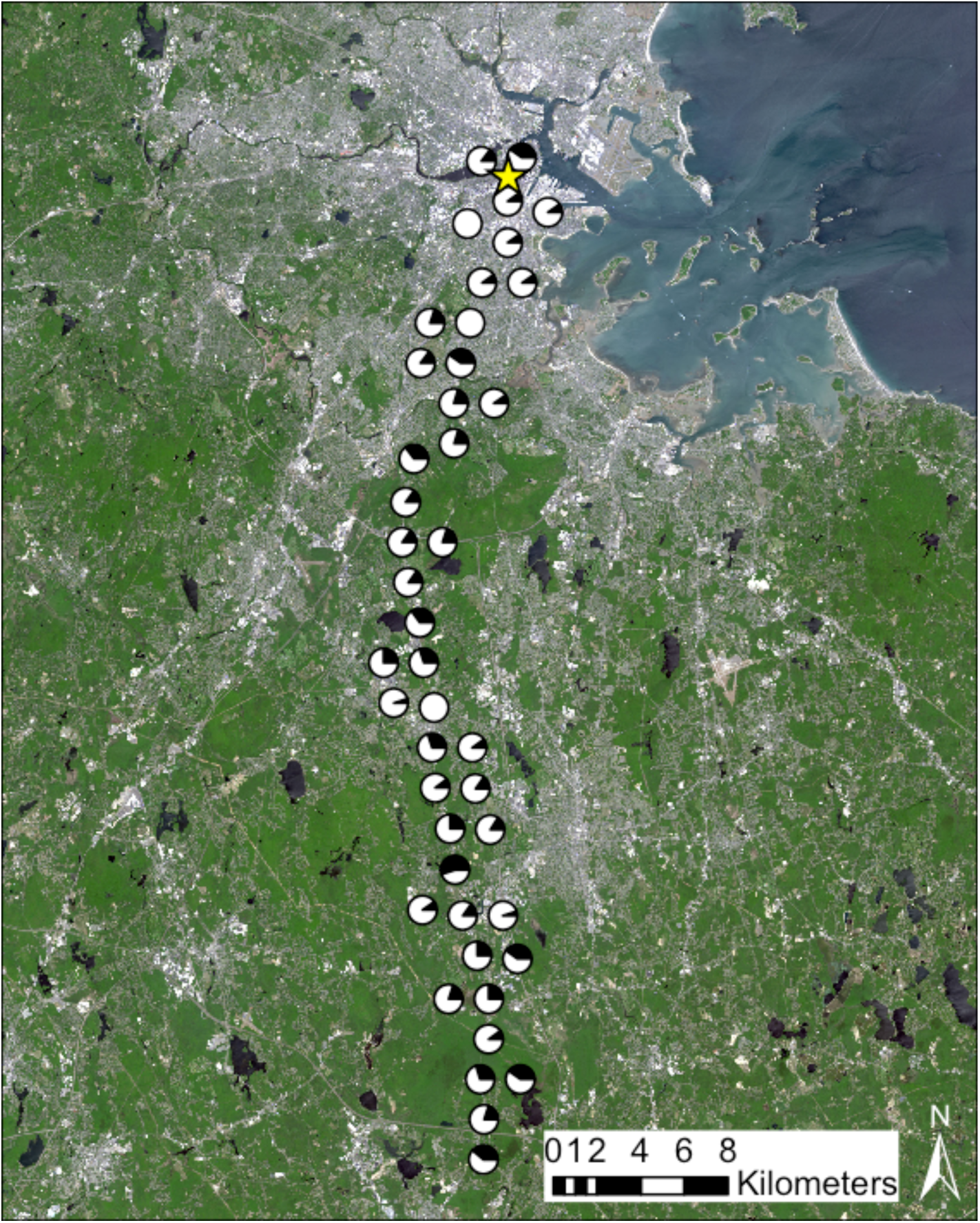
Map depicting the location and HCN frequency in populations sampled in Boston, USA (*n* = 43). The proportion of black colouration in pie charts reflects the frequency of HCN in that population. Pie chart locations were jittered to avoid overlap. We defined the urban center of Boston as Boston Common, indicated by the star symbol. The image is from the Landsat 8 satellite and was captured on 24 May 2015.

**Fig. S6.**
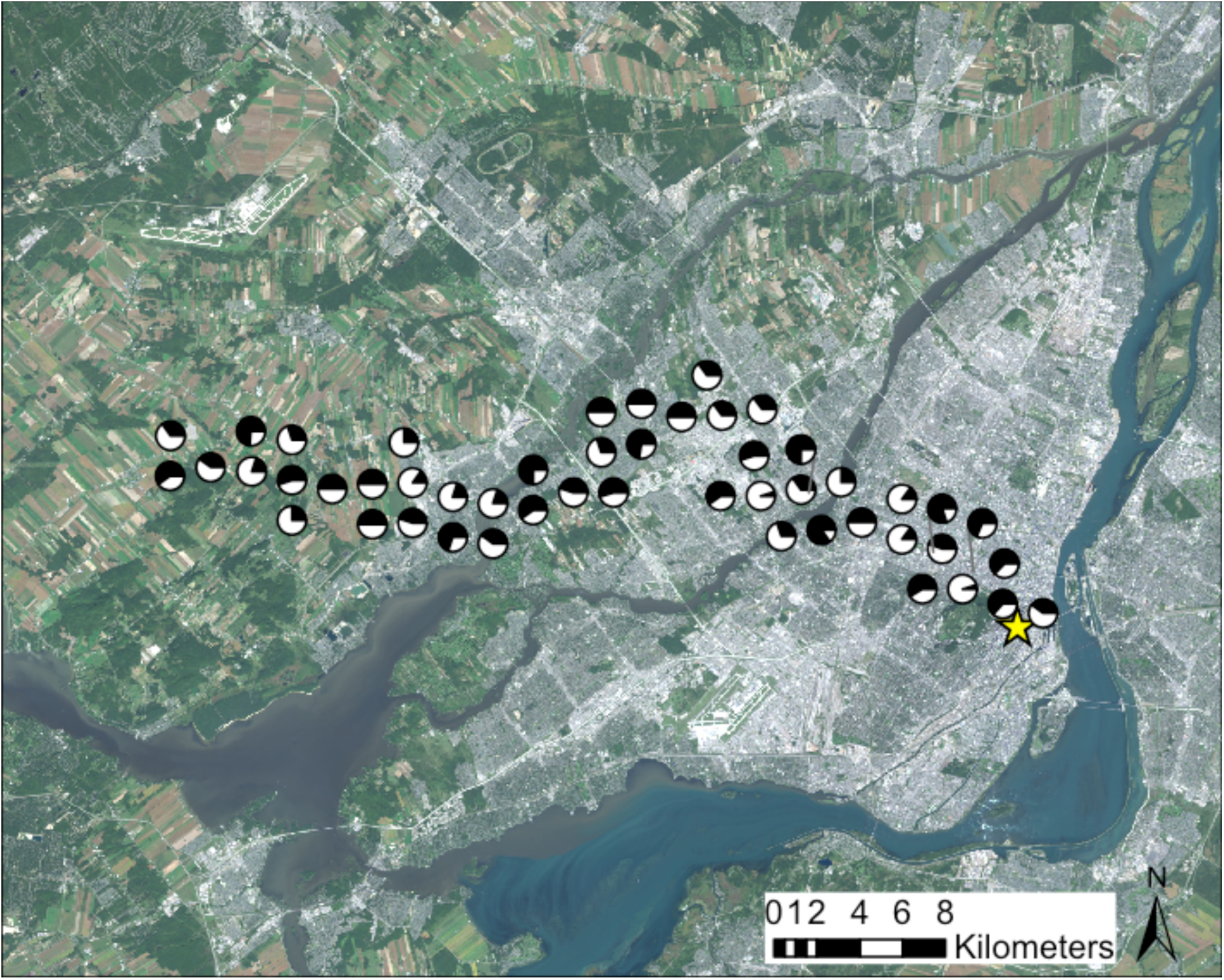
Map depicting the location and HCN frequency in populations sampled in Montreal, Canada (*n* = 49). The proportion of black colouration in pie charts reflects the frequency of HCN in that population. Pie chart locations were jittered to avoid overlap. We defined the urban center of Montreal as the intersection of Boulevard Robert-Bourassa and Boulevard René-Lévesque Ouest, indicated by the star symbol. The image is from the Landsat 8 satellite and was captured on 18 September 2015.

**Fig. S7.**
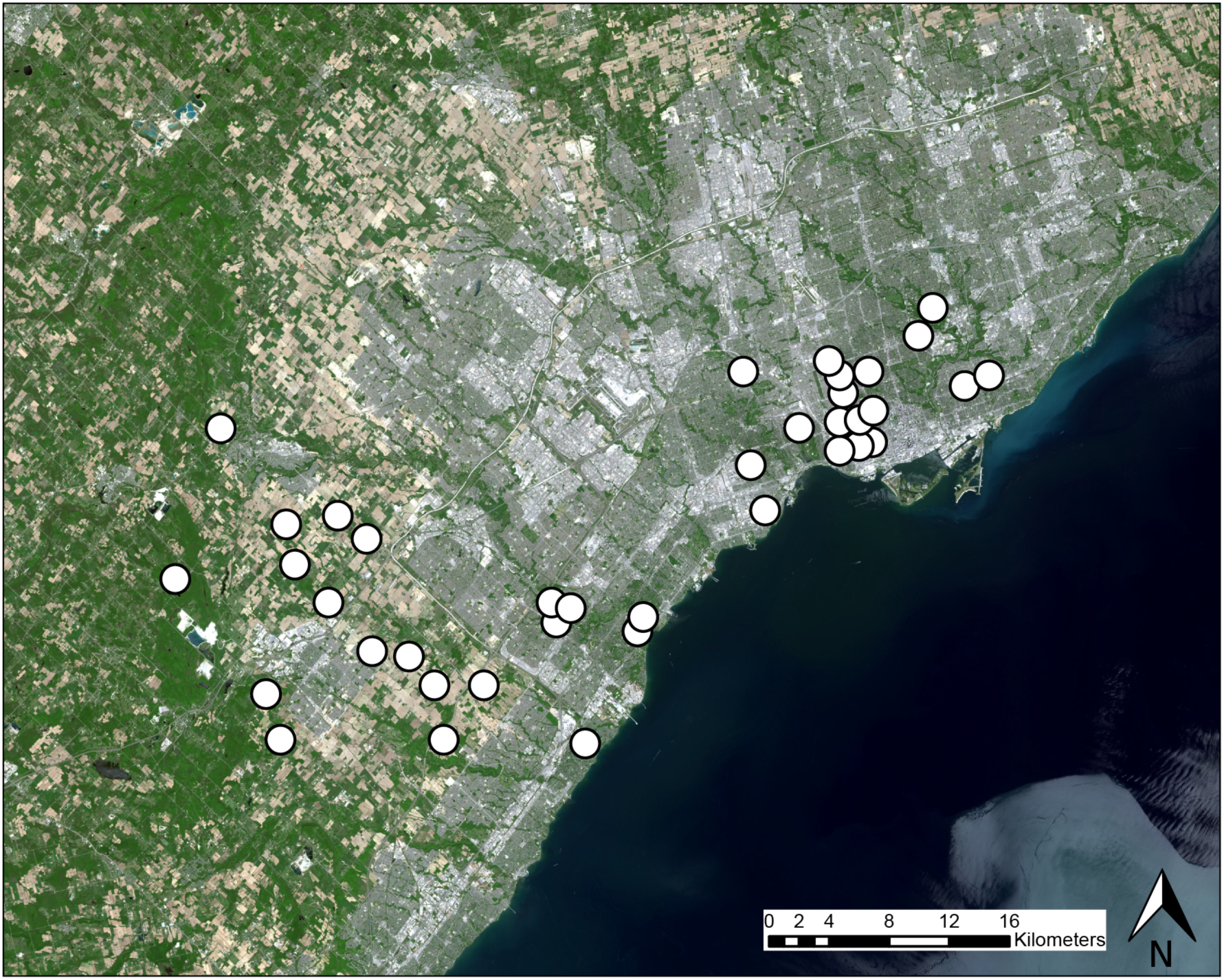
Map showing the location of experimental clover populations in our field experiment in the Greater Toronto Area during summer 2015 (*n* = 38). The image is from the Landsat 8 satellite and was captured on 31 May 2014.

**Fig. S8.**
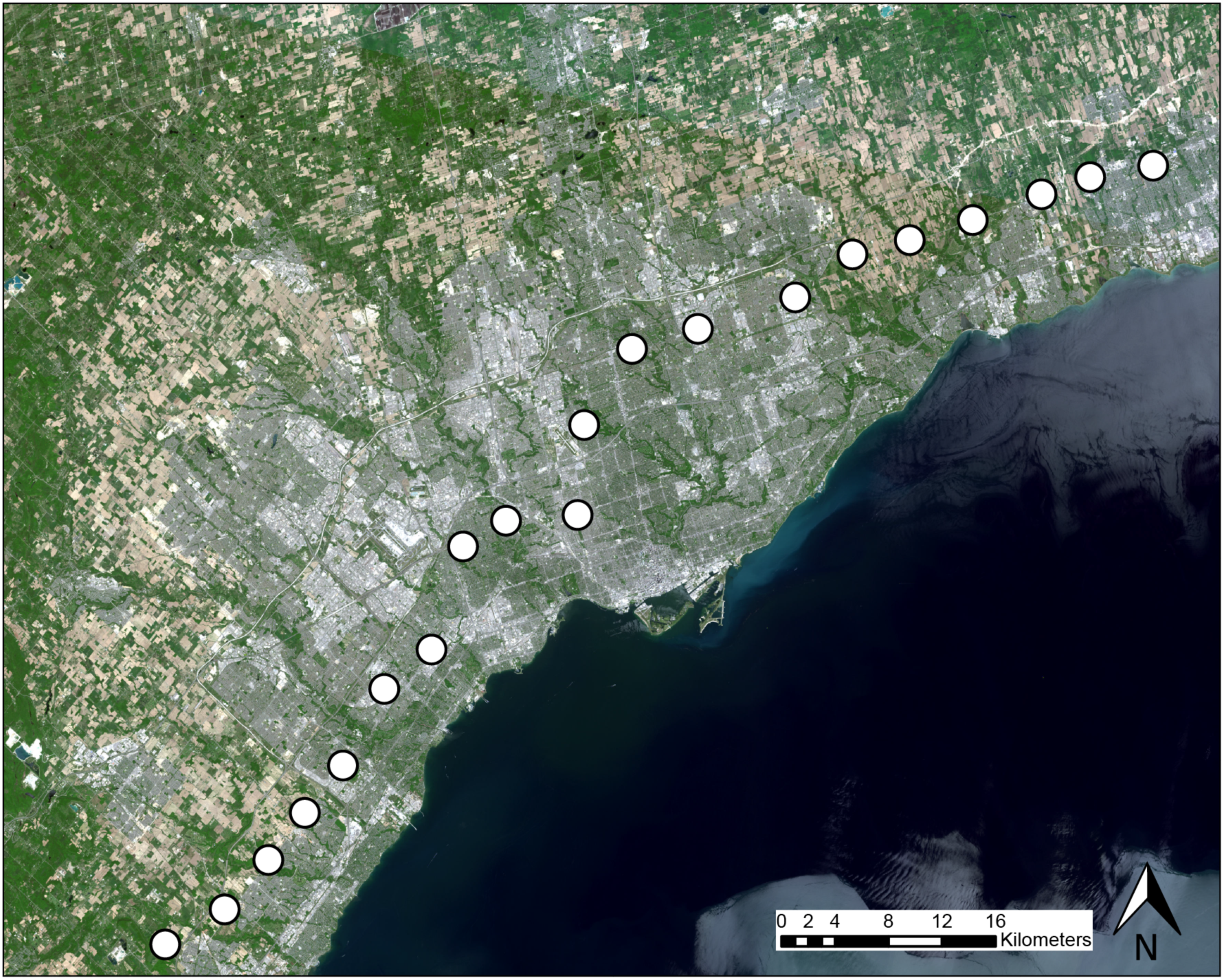
Map showing the locations of iButton temperature probes that were dispatched across the Greater Toronto Area during winter 2015 (*n* = 20). iButtons recorded ground temperature measurements every hour from 28 Jan to 27 Mar 2015. The image is from the Landsat 8 satellite and was captured on 31 May 2014.

**Fig. S9.**
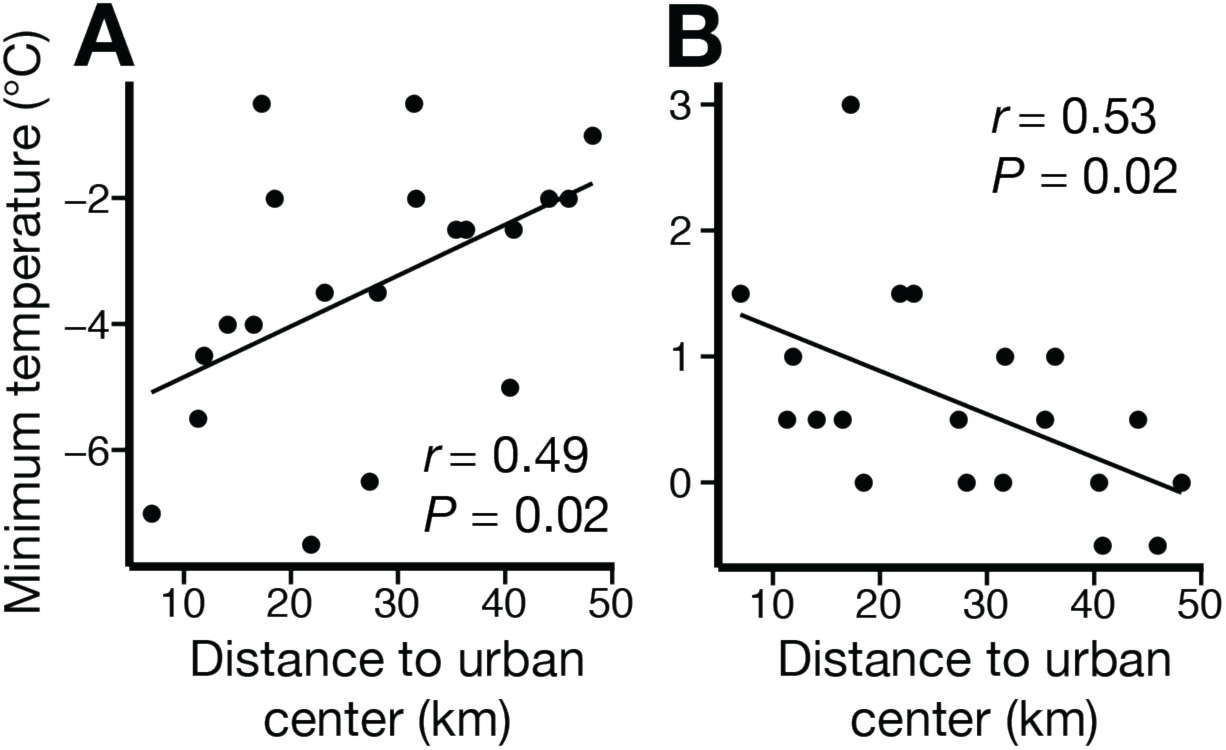
Representative images of: (**A**) positive and (**B**) negative relations between minimum temperature and distance from the urban center. The ‘relative urban coldness’ index values for the left and right panels are 0.081 and –0.034, respectively, which was equivalent to the slope. When the index is positive, temperatures are colder in the city than outside of the city (the winter urban cold island effect) and when the index is negative, temperatures are warmer inside the city than outside of the city (the urban heat island effect). Minimum temperature values were taken from iButton temperature probe data. The data in panel (**A**) were captured on 4 March 2015, and the data in panel (**B**) were captured on 26 March 2015. Statistics were calculated using linear regression.

**Fig. S10.**
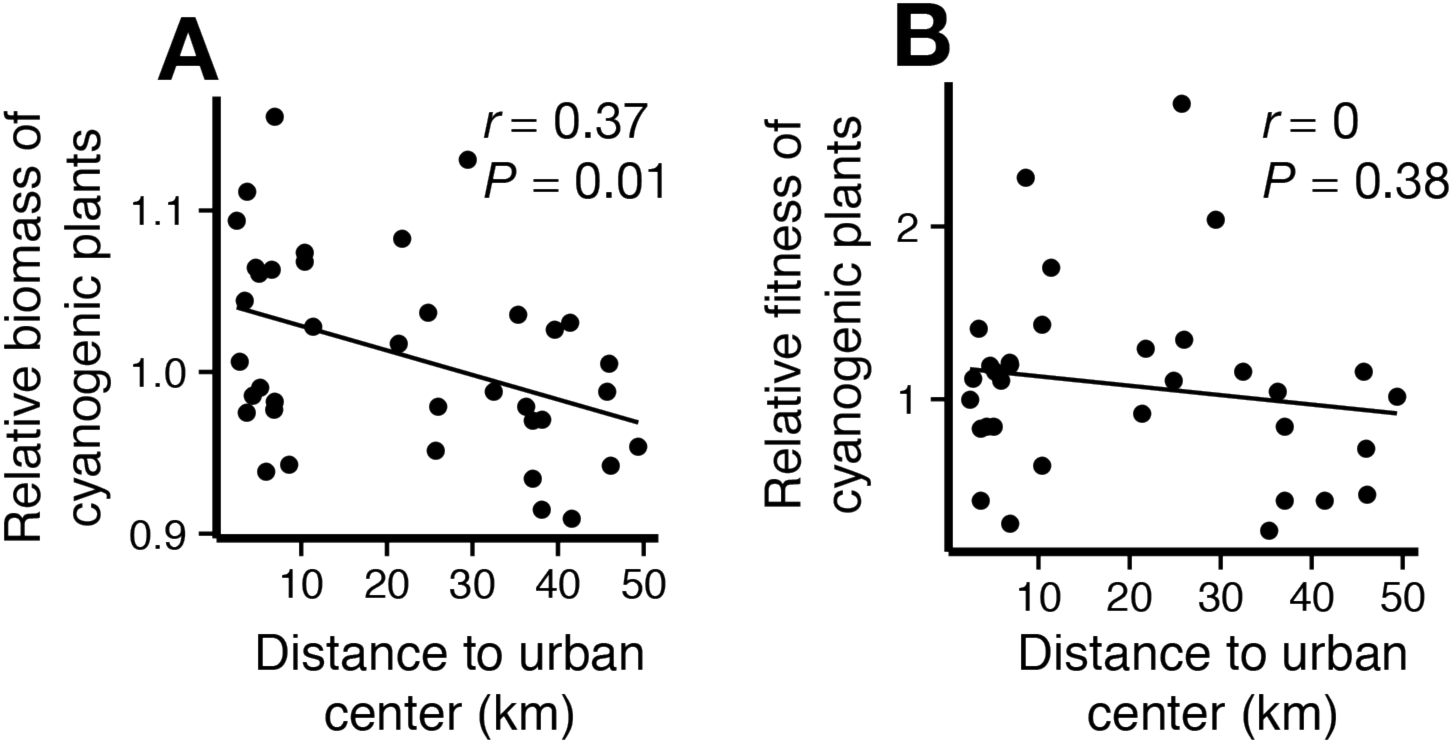
Selection on cyanogenesis during the field experiment in summer 2015. (**A**) Relative to cyanogenic plants, cyanogenic plants had higher final biomass in urban areas and lower final biomass in non-urban areas. We interpreted this result as evidence of natural selection for cyanogenesis in the urban populations. This result contrasts the results of our natural population surveys. Whereas we observed selection operating through final biomass, (**B**) there was no selection on cyanogenesis via maternal fitness (seed mass) across the urbanization gradient. Statistics were calculated using linear regression.

## Supplementary information

<u>Supplementary text S1: *Study System*</u>

*Trifolium repens* L. (Fabaceae), white clover, is a globally distributed perennial plant common to lawns and disturbed areas. Native to Eurasia, white clover has spread globally because of human introduction as a nitrogen-fixing pasture crop and soil stabilizer (Baker and Williams 1987). Self-fertilization is rare in *T. repens* due to a gametophytic self-incompatibility system (Douglas and Connolly 1989). While individual plants produce many seeds, seed recruitment in wild populations is rare (Barrett and Silander 1992). Clover is highly clonal, and produces trailing stolons that root at the nodes (Kemball and Marshall 1995).

In wild populations of white clover, some plants produce hydrogen cyanide (HCN)—a general toxin that inhibits enzymes containing heavy metals, including cytochromes in the electron transport chain (Nahrstedt 1985; Leavesley et al. 2008). The cyanogenesis polymorphism in *T. repens* has been studied for over a century (Mirande 1912; Armstrong et al. 1913) and is regulated by alleles at two independently segregating diallelic loci—*CYP79D15* and *Li. CYP79D15* (hereafter *Ac*) regulates the initial step of cyanogenic glycoside biosynthesis, and *Li* encodes the enzyme linamarase (Olsen et al. 2007, 2008). Cyanogenic glycosides (linamarin and lotaustralin) are stored in vacuoles (Poulton 1990), and linamarase is stored in the cell wall (Kakes 1985). Following tissue disruption, these two components are brought into contact to release free HCN. A single copy of the dominant allele at both loci (*Ac* and *Li*) is required for a plant to produce HCN (cyanotype *AcLi*). Plants without one or both dominant alleles are acyanogenic (cyanotypes *Acli*, *acLi*, and *acli*). Dominant alleles confer the presence of the corresponding nucleotide sequence, and recessive alleles confer the compete absence of the genes due to complete deletions at *Ac* and *Li* (Olsen et al. 2013).

Early work examining the distribution of cyanogenesis in wild populations of *T. repens* found that cyanogenesis decreases in frequency toward high latitudes (Daday 1954b) and elevations (Daday 1954c). Many authors have since reported similar patterns (de Araújo 1976; Brighton and Horne 1977; Till 1987; Till-Bottraud et al. 1988; Ganders 1990; Caradus and Forde 1996; Pederson et al. 1996; Richards and Fletcher 2002; Majumdar et al. 2004; Kooyers and Olsen 2013; Oliveira et al. 2013; Shrestha Malla 2014), and several have explored their mechanistic basis. In warm climates, cyanogenic plants are hypothesized to outperform acyanogenic plants because of increased herbivore pressure and aridity. In addition to the present study, several studies have shown that cyanogenic plants are less damaged by herbivores than acyanogenic plants (Angseesing 1974, Dirzo and Harper 1982a; Horrill and Richards 1986; Kakes 1989; Pederson and Brink 1998; Saucy et al. 1999; Thompson and Johnson 2016), but the evidence for differential tolerance to aridity is mixed (Foulds and Grime 1972; Kooyers et al. 2014).

Cold winter climates are hypothesized to select against cyanogenesis because cyanogenic plants experience self-toxicity when free HCN is released following freezing-induced tissue damage (Stout et al. 1980, 1986; Gleadow and Møller 2014). Several studies have shown that cyanogenic *T. repens* plants are more damaged by freezing temperatures than acyanogenic plants (Daday 1958, Dirzo and Harper 1982b; Caradus and Christie 1998; Olsen and Ungerer 2008), and artificial selection for frost hardiness in *T. repens* causes selection against cyanogenesis (Caradus and Eerens 1989). Perhaps to avoid frost damage, *T. repens* decreases the production of cyanogenesis components when temperatures drop below 8°C (Collinge and Hughes 1982). Physiological studies in *Lotus corniculatus*, another legume with a cyanogenesis polymorphism, demonstrated that cyanogenesis irreversibly inhibits leaf respiration following exposure to freezing temperatures (Brighton and Horne 1977). Additional studies have found that freezing causes HCN release in live tissue, and that highly cyanogenic plants across multiple species have reduced freezing tolerance (Lebeau 1966; Stout et al. 1981; Kuti and Konoru 2006).

<u>*Supplementary text S2: Opposing natural selection on cyanogenesis across seasons*</u>

We observed significant natural selection favouring cyanogenesis in cities during the summer (*β*= −1.5 × 10^−3^ ± 5.6 × 10^−4^, *F*_1, 36_ = 7.24, *P* = 0.01) (Fig. S10A). Relative to acyanogenic plants, the final biomass of cyanogenic plants was larger in urban populations and smaller in non-urban populations. This pattern only held for *AcLi* genotypes; relative fitness did not change across an urbanization gradient for *Acli* (*β* = 1.0 × 10^−4^ ± 1.4 × 10^−2^, *F*_1, 36_ = 0.005, *P* = 0.94) or *acLi* (*β* = 6.1 × 10^−4^ ± 1.3 × 10^−3^, *F*_1, 36_ = 0.22, *P* = 0.64) genotypes, indicating that the cyanogenic phenotype *per se* is responsible for the pattern. This observation is consistent with our original prediction that cyanogenesis would be favoured in cities. Recent experimental studies have linked cyanogenesis with tolerance to dry soils and hypothesize that water stress may be partially responsible for the evolution of clines in cyanogenesis (Kooyers et al. 2014). While we did not quantify soil moisture during our experiment, urban populations may have experienced increased water stress because of warmer temperatures and increased impervious surface cover (Whitlow et al. 1992). One possible problem with this hypothesis is that previous studies have linked dominant alleles at *CYP79D15* to water stress tolerance (Kooyers et al. 2014), but we did not find a fitness advantage of *Acli* genotypes. We note that selection was only acting through final biomass, and not lifetime biomass or sexual fitness (Fig. S10).

<u>*Supplementary text S3: Alternative explanations for cyanogenesis clines along urbanization gradients*</u>

S3.1—*Mowing and fertilization experiment*

S3.1-1—*Experimental design*

Urbanization can be associated with clines in human activities such as lawnmower use and fertilizer application, both of which have the potential to differentially affect cyanogenic and acyanogenic plants. We tested whether a differential response to mowing and fertilizer could explain the observed urban-rural cyanogenesis clines using a growth chamber experiment. We took four stolon cuttings from each of 89 plants for a total of 356 plants. Cyanotypes of these 89 plants were: 35 *AcLi*, 20 *Acli*, 14 *acLi*, and 20 *acli*. Cyanotypes were determined as in the field experiment described above. All plants were grown from seed collected from different genotypes in the experiment described by Thompson and Johnson (2016). All cuttings weighed between 0.5 g and 0.8 g. We replaced cuttings that did not survive transplanting within the first four days of the experiment. Two plants died after this period and thus our final analysis consisted of 354 plants. Cuttings were propagated into 500 mL pots (STD0400, ITML, Brantford, ON) filled with soil (Sunshine Mix #2, Sun Gro Horticulture, Agawam, USA). Pots were randomized into a growth chamber with day:night temperatures of 25:22 °C, with 16 hours of 500 *μ*mol/m^2^/s light. All pots were contained in small plastic cups to prevent mixing of water between pots, which were held in solid plastic trays.

We applied damage and fertilization treatments in a 2 × 2 fully factorial design. The damage treatment consisted of undamaged control plants and plants that were damaged in a manner that simulated lawnmower damage on lawns. We housed plants beneath a wooden plant with a hole in the center that was level with the upper rim of the pots, and removed all tissue that was greater than 1.5 cm above the upper rim of pots. Damage treatments were applied three times, on the 20^th^, 30^th^, and 40^th^ day of the experiment. Our fertilizer treatment included control plants (no fertilizer) and fertilized plants, which received 1.5 g of slow-release fertilizer at the beginning of the experiment (Nutricote Total 13-13-13 With Minor Nutrients, Florikan, Sarasota, FL). The experiment was terminated on day 58, at which time we harvested the aboveground biomass, dried the tissue for 72 hr at 60 °C and then weighed the tissue to the nearest 0.1 g.

The ability for plants to regrow following damage represents an estimate of tolerance. Because of issues with interpreting tolerance estimates when multiplicative and additive data are used (Wise and Carr 2008), we conducted our analysis using log-transformed data because the amount of tissue removed was related to the size of plants. Our conclusions about whether HCN influences tolerance do not change if untransformed data are used.

S3.1-2—*Statistical analysis*

Because damage and fertilization treatments influenced plant biomass to very different degrees, we analyzed the main effects of each in a mixed-effects one-way ANOVA with one treatment as a fixed categorical variable, and genotype and the other treatment as random effects. We used a mixed-effects three-way ANOVA to analyze whether cyanogenesis influenced the response to damage or fertilization. Our full statistical model was:

*W = intercept + damage + fertilization + HCN + damage × fertilization + damage × HCN + fertilization × HCN + damage × fertilization × HCN + genotype + error*

where *W* is the log-transformed fitness component (final vegetative biomass or lifetime floral biomass), damage and fertilization were factors representing experimental treatments, HCN is the presence (cyanogenic) or absence (acyanogenic) of HCN, and genotype was a categorical random effect.

S3.1-3—*Results*

Mowing and fertilization both had significant effects on fitness components. In univariate mixed-effects models, damaged plants had approximately 20% lower final vegetative biomass (*F*_1, 264_ = 64, *P* < 0.01) and produced 20% less lifetime sexual biomass than undamaged plants (*F*_1, 264_ = 36, *P* < 0.01). Similarly, unfertilized plants had approximately 52% lower final vegetative biomass (*F*_1, 264_ = 692, *P* < 0.01) and produced 72% less lifetime sexual biomass (*F*_1, 264_ = 995, *P* < 0.01) than fertilized plants. There was no difference in vegetative biomass between cyanogenic and acyanogenic plants (*P* > 0.5), but acyanogenic plants produced 18% more sexual biomass than cyanogenic plants (*F*_1, 87_ = 5.06, *P* = 0.03).

Cyanogenesis and fertilization did not influence the ability of plants to tolerate simulated mowing treatments (all interactions: *P* > 0.20), and cyanogenesis did not influence plant response to fertilization (*P* = 0.73). That is, cyanogenesis did not influence plant fitness in response to any treatment or treatment combination. We conclude that differential tolerance to mowing and/or fertilization does not cause clines in cyanogenesis along urbanization gradients.

S3.2.2—*Salinity and clines in cyanogenesis*

Recent studies have shown that soil salinity can upregulate HCN production in *T. repens* (Ballhorn and Elias 2014). Several additional studies have shown that soil salinity is higher in urban areas relative to non-urban areas (Cunningham et al. 2008). Given that highly cyanogenic plants are the most susceptible to freezing damage (Caradus et al. 1989), urban plants may be more highly damaged from freezing than non-urban plants because of higher HCN concentration in their tissue. This would be exaggerated under a winter urban cold island scenario. The link between soil salinity, cyanogenesis, and freezing tolerance in *T. repens* requires further investigation.

S3.2.3—*Aridity and cyanogenesis clines*

Clines in cyanogenesis are often associated with aridity, with higher frequencies of cyanogenic plants in arid habitats (Kooyers and Olsen 2013). Arid soils are often associated with urban environments (Whitlow et al. 1992), and so if aridity was responsible for cyanogenesis clines then we expect cyanogenesis to be most frequent in cities. While our field experiment is consistent with natural selection favouring cyanogenesis in the summer in urban habitats (Fig. S10A), if aridity were the primary driver of cyanogenesis cline evolution, the clines would likely be in the opposite direction than we observed (i.e., higher frequency of cyanogenesis in urban environments). Given that this is not the case we conclude that aridity is unlikely to explain the observed clines.

<u>S4. Additional acknowledgements</u>

We are grateful the following individuals who made their properties available for the field experiment conducted herein: Lee Adamson, Spencer Barrett&Tracy Keenan, Gordan Bergman, Paula Brown, Sheila Boonstra, Lucy Cofini, Bill&Mary Cole, Sheila Colla, Nick Collins&Megan Pallett, Nettie Cronish&Jim Urquhart, Tim Dickinson, Susan Dixon, Phil Rudz, Mark Engstrom, Elias&Mike, Bruce&Ann Falls, Benjamin Gilbert&Kate Kirby, Corey Goldman, Tess Grainger, Darryl Gwynne, Christina Iwaschko&Kristin Brink, Laura Junker, Aaron LeBlanc&Audrey Reid, Tracy&Ted Lee, Janna Levitt, Shannon McCauley&Stephan Sneider, Nina Michalak, Gayle Redshaw, Fiona Reid, Christoph Richter, Dan&Shelby Riskin, Chris Searcy&Michelle Afkhami, Gary&Diane Sprules, Bev Sutton, and Janet Trinder.

